# High throughput gene expression profiling of yeast colonies with microgel-culture Drop-seq

**DOI:** 10.1101/416966

**Authors:** Leqian Liu, Chiraj Dalal, Ben Heineike, Adam Abate

**Author notes:** Correspondence and requests for materials should be addressed to A.R.A.

## Abstract

Yeasts can be engineered into “living foundries” for non-natural chemical production by reprogramming their genome using a synthetic biology “design-build-test” cycle. While methods for “design” and “build” are scalable and efficient, “test” remains a labor-intensive bottleneck, limiting the effectiveness of the genetic reprogramming results. Here we describe Isogenic Colony Sequencing (ICO-seq), a massively-parallel strategy to assess the gene expression, and thus engineered pathway efficacy, of large numbers of genetically distinct yeast colonies. We use the approach to characterize opaque-white switching in 658 *C. albicans* colonies. By profiling transcriptomes of 1642 engineered *S. cerevisiae* strains, we use it to assess gene expression heterogeneity in a protein mutagenesis library. Our approach will accelerate synthetic biology by allowing facile and cost-effective transcriptional profiling of large numbers of genetically distinct yeast strains.

## Introduction

The baker’s yeast *Saccharomyces cerevisiae* is easy to culture, non-infectious to humans, and has powerful tools for modifying its genome (1–3). For these and other reasons, *S. cerevisiae* has been engineered into “living foundries” that synthesize high-value chemicals from renewable feedstock (4,5). To maximize production, the cells must be genetically engineered to convert feedstock into product. Optimally engineering a cell’s genome is a challenging process that often relies on a “design-build-test” cycle, in which variants are constructed and tested for the desired activity (6,7). The effectiveness of this process depends on the number of strains that can be built and tested. While advances in computation, DNA synthesis, and genome editing enable facile construction of large libraries (>10), testing these strains remains a costly bottleneck as it requires each strain to be individually isolated, cultured, and assayed for activity (6).

Droplet microfluidics has markedly increased the throughput of strain testing. Similar to flow cytometry, droplet microfluidics analyzes individual cells at kilohertz rates; however, a unique advantage of droplets is that they allow cells to be characterized based on phenotypes not detectable with flow cytometry, like extracellular analyte consumption, product secretion, and interactions with other cells (6,8–10). Nevertheless, a limitation of the approach is that it usually requires a fluorogenic assay, which is not possible for many phenotypes and can be challenging to optimize for the requisite water-in-oil emulsions (11,12). Moreover, these assays are usually specific to a given target, making it difficult to extend the approach to many targets.

Gene expression profiling by mRNA sequencing is a standard approach for characterizing cell phenotypes and has been used for applications like characterizing the cell cycle, transcriptional rewiring, pathway efficiency assessment, and metabolic flux analysis (13–16). Moreover, new droplet microfluidic based single cell RNA-seq (scRNA-seq) has demonstrated gene expression profiling of tens of thousands of single cells per experiment (17–19). However, these approaches have been optimized for mammalian cells, which are larger and contain more RNA than the yeast cells commonly used as living foundries. Consequently, droplet microfluidic scRNA-seq is ineffective when applied to fungal and other microbial cells. While a recent study has demonstrated single yeast cell sequencing with Fluidigm C1 platform (20), the throughput is limited and not adequate for testing strains from libraries. Additionally, scRNA-seq is subject to significant biological and technical noise due to the dynamic nature of gene expression and the tiny amount of RNA available for sequencing. To enable effective, general, and scalable strain testing, a new approach for characterizing gene expression of engineered strains is needed that has the generality of single cell RNA sequencing and is applicable to fungal cells.

In this paper, we present Isogenic Colony Sequencing (ICO-seq), a general approach for profiling the gene expression of cultivable cells at high throughput. The key innovation of ICO-seq is the coupling of hydrogel droplet culture of single cells with barcoded RNA sequencing (Drop-seq). Hydrogel culturing amplifies single cells into an isogenic colony of tens to hundreds of cells; this provides ample RNA for deep sequencing of the colony and reduces error due to noisy single cell gene expression profiles. The resultant profiles correspond to individual strains and can be used to screen for the desired phenotype. To demonstrate and validate ICO-seq, we use it to study white-opaque switching in *Candida albicans* and to assess expression heterogeneity of a *Saccharomyces cerevisiae* ARO4 regulatory domain mutagenesis library. We anticipate our approach will aid in the generation of optimized strains by allowing rich gene expression information to be collected from tens of thousands of genetically distinct fungal strains. This platform can be readily implemented into the “design-build-test” cycle of synthetic biology.

## Results

### ICO-seq Workflow

The goal of ICO-seq is to obtain high-quality gene expression profiles from genetically and epigenetically distinct yeast colonies strains with the scalability of droplet approaches. To enable this, ICO-seq integrates two droplet microfluidic technologies: the encapsulation and culture of single cells in hydrogel beads (21) and Drop-seq RNA sequencing (17). In the cultivation step, single yeast cells are encapsulated in 90 μm agarose beads and immersed in culture medium; this expands a single cell into an encapsulated colony of hundreds of isogenic clones, averaging single cell noise and amplifying the amount of RNA available for barcoded sequencing (Figure 1a). In the barcoding step, the hydrogel cultures are paired with barcoded Drop-seq beads and encapsulated in droplets using a custom microfluidic device; lysis buffer is also included, lysing the cells and releasing mRNA for oligo(dT) capture and barcoding, as in standard Drop-seq protocols (Figure 1b). The resultant barcoded data is processed similarly to conventional single cell RNA sequencing data, first grouped by barcode and then subjected to gene expression analysis (Figure 1c). The principal difference between ICO-seq and Drop-seq is that barcode groups correspond to isogenic colonies of many of cells, rather than single cells.

**Figure 1.**
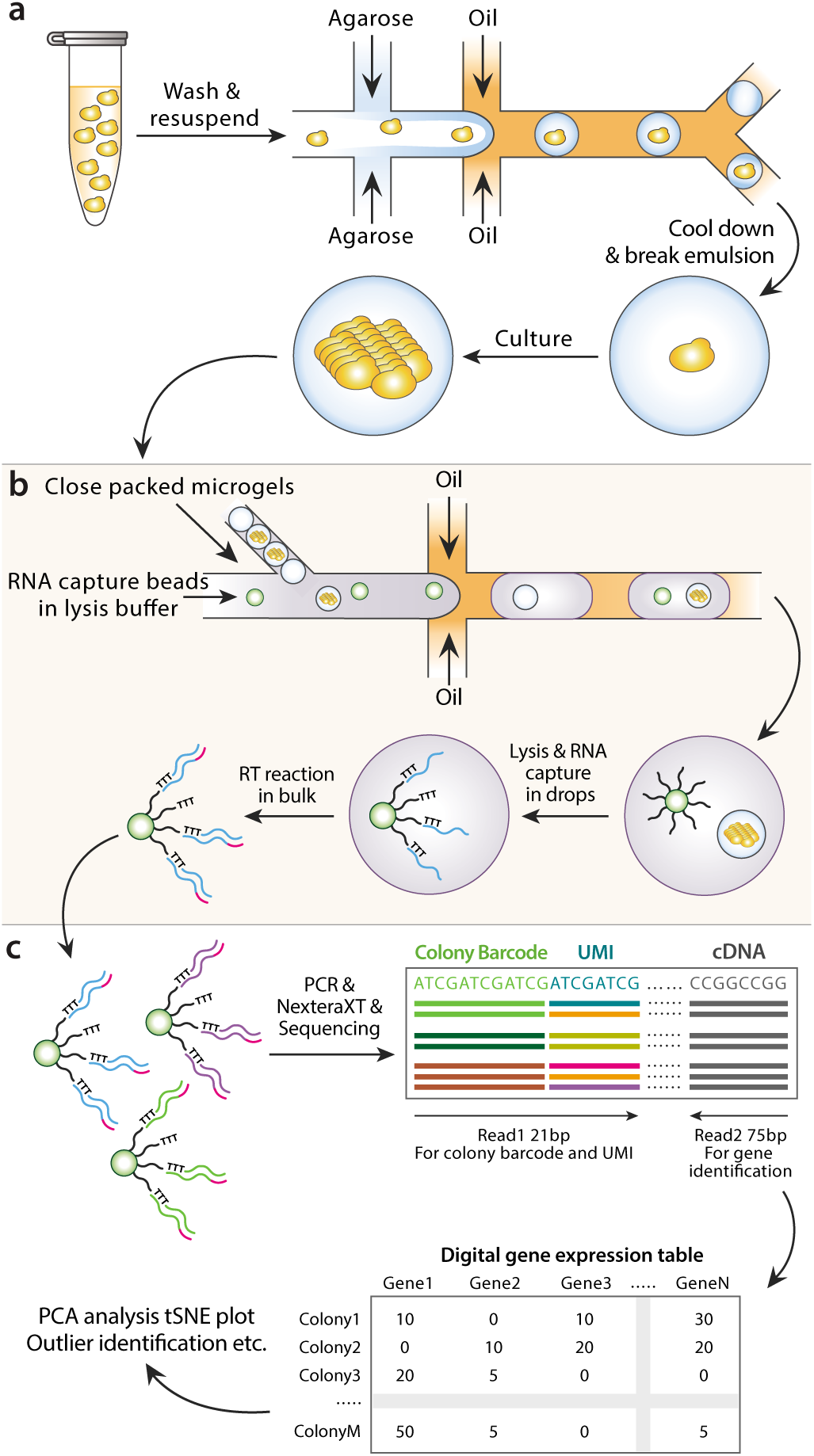
Schematic of ICO-seq workflow. (a) Colony-containing agarose microgels are generated by encapsulating single yeast cells into agarose microdroplets followed by microgel recovery and colony formation within the microgel. (b) mRNA is converted to cDNA from single colonies by co-encapsulation of microgels and barcoded RNA capture beads followed by in-droplet hybridization and a bulk reverse transcription reaction after droplets are demulsified. (c) Sequencing libraries are prepared from cDNA containing barcoded RNA capture beads following the Drop-seq protocol. Following sequencing, digital gene expression tables for individual yeast colonies are generated and analyzed.

### Microfluidic design and operation in ICO-seq

ICO-seq utilizes microfluidic technologies for encapsulating single cells in microgels and pairing the resultant isogenic cultures with barcoded beads for sequencing. The hydrogel matrix must be carefully selected to allow microfluidic synthesis of the microgels, single cell culture, and Drop-seq. Agarose is a suitable hydrogel material for many biological applications and is compatible with microfluidic microgel fabrication and Drop-seq (21–23). A key property of agarose is that it melts at moderate temperatures (~90°C), allowing it to flow through microfluidic channels and be formed into liquid droplets. Upon cooling, the droplets solidify into elastic microgels. If cells are included in the droplets, they are embedded in the resultant microgels (21,23). To utilize agarose in this way, we thus require a microfluidic device that can emulsify the molten solution into monodispersed droplets. Because the elevated temperature (~50 °C) required to keep molten agarose from gelling may harm cells, the melt and cell suspensions are introduced via separate inlets using a “co-flow” geometry (Figure 2a, yellow box). Due to low Reynolds number laminar flow, these miscible solutions merge but do not mix until encapsulated in droplets in the flow-focus junction. In addition, the droplets cool down rapidly due to their large surface area to volume ratio.

**Figure 2.**
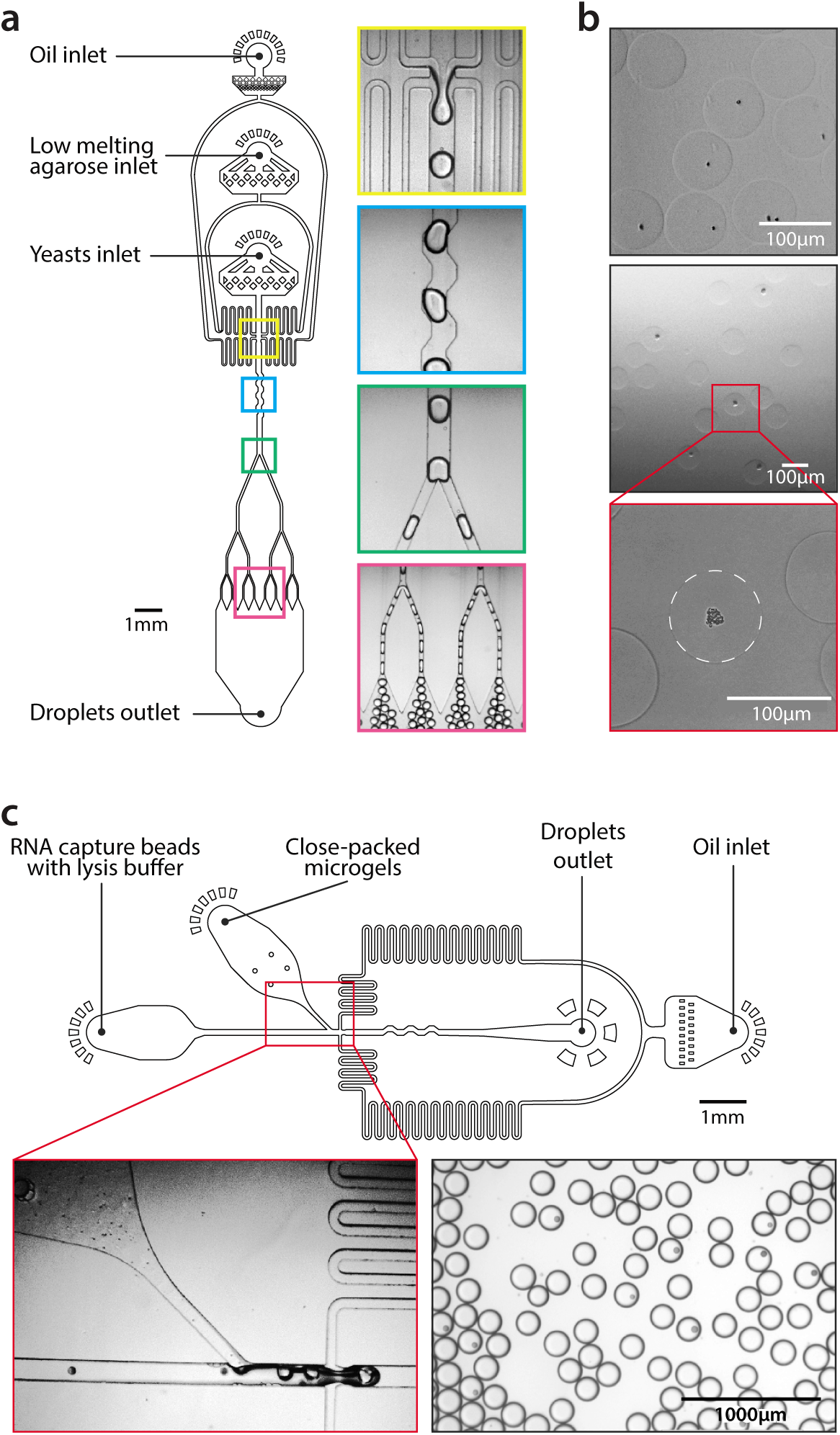
ICO-seq microfluidic device design and operation. (a) Microfluidic device design for high throughput single yeast encapsulation into agarose microgels. From top to bottom, images showing droplet generator, droplet mixer, the first droplet splitter and the third droplet splitter. (b) Yeast colony formation within agarose microgels. Top: single cell encapsulated agarose microgels; middle and bottom: colony containing agarose microgels. (c) Microfluidic device design for co-encapsulation of agarose microgels and barcoded RNA capture beads. Images on the bottom showing co-encapsulation step (left panel) and droplets generated with this device (right panel).

For ICO-seq to be useful, it must be scalable to tens of thousands of cells, which necessitates rapid encapsulation of cells in the microgel spheres. However, molten agarose is viscous, slowing droplet generation. To increase droplet production rate, we thus employ sequential droplet splitting (24). A large droplet (3200 pL) is formed at high flow rates and subsequently bisected into eight sister droplets (400 pL) (Figure 2a, green and red boxes). The net throughput of this device (5 KHz) is ~8-fold faster than a single droplet generator creating 400 pL droplets directly. To ensure the daughter droplets contain an equivalent composition of cell and agarose solution, a mixing channel stirs the large droplets before they reach the first split (Figure 2a, blue box).

The molten agarose droplets are collected on ice to solidify, then the oil is removed via demulsification and the droplets are washed and transferred to an aqueous solution. To ensure a fraction of the microgels contain single yeast cells, the cell suspension is diluted such that ~30% of droplets contain one cell in accordance with a Poisson distribution (Figure 2b, top panel). This cell encapsulation rate yields ~10% of microgels containing cultured yeast colonies, since some of the encapsulated cells do not grow and form colonies. The washed microgels are then transferred into culture media. The porous agarose allows the media to perfuse through the gels, allowing the cells to grow to form colonies (Figure 2b, middle and bottom panels). The colony size can be tuned by controlling the culture properties, including the media composition, and duration and temperature (21,23).

The second step in ICO-seq is to capture and barcode each microgel culture’s mRNA. This requires lysing the cultures in a droplet containing a barcoded oligo(dT) capture bead, following the Drop-seq protocol (17). It thus requires a microfluidic device that encapsulates each microgel culture with a barcoding bead (Figure 2c). Prior to injection into the device, the microgels are close-packed by centrifugation. Due to the monodispersity of the microgels and their close packing, they flow regularly into the droplet generator, allowing efficient loading (25). Barcoded RNA capture beads are suspended in lysis buffer at a concentration such that on average 10% of the droplets contain a single bead, and are introduced via a parallel inlet, spacing the microgels just before droplet generation. Upon encapsulation, lysis buffer diffuses into the microgels. This releases polyadenylated mRNA from the colony so that it hybridizes to the oligo(dT) barcodes on the capture beads. This process is repeated for hundreds of thousands of droplets in parallel. For each colony that is co-encapsulated with a capture bead, its mRNA is captured onto the uniquely barcoded oligo(dT) capture probes on the bead surface. The droplets are then demulsified, and the recovered beads are subjected to the remaining steps of Drop-seq, including reverse transcription (Figure 1b), PCR, and tagmentation for Illumina sequencing library preparation. The library is sequenced and the resulting gene expression data is analyzed (Figure 1c).

### Analysis of white-opaque switching in *C. albicans* using ICO-seq

*C. albicans* is an opportunistic commensal yeast that can colonize and invade human tissue when the human immune system is compromised or the competing microbial flora eliminated (26). It has been hypothesized that the ability of *C. albicans* cells to colonize warm-blooded hosts is enhanced by its ability to switch between multiple cell types, amongst which are the white and opaque cell types (27,28). White-opaque switching results in two distinct types of cells from the same genome that vary significantly from each other in size, shape, susceptibility to host defense and mating-competence (29). Perhaps most interesting to us, white-opaque switching is heritable; in other words, when an opaque cell switches to white, the resulting progeny continue to be white (it is important to note that these white cells can switch back to opaque cells, albeit at a low frequency).

Here we use an opaque diploid *C. albicans* strain engineered such that YFP (yeast fluorescent protein) replaces one copy of the WH11 gene. Since the WH11 promoter is active only in white cells(30) and its expression correlates with the commitment of cells being white, YFP expression under WH11 control monitors the switch from opaque to white (29). With ICO-seq, which provides gene expression profiles of isogenic colonies, the presence of YFP mRNA can be used to define colony type. In this experiment, we encapsulated opaque *C. albicans* cells in agarose microgels followed by colony formation through cultivation. Due to switching from opaque-to-white cells, opaque and white *C. albicans* colonies (as well as some mixed cultures) existed at the end of cell culturing. These colonies were analyzed for gene expression using ICO-seq (658 colonies with at least 600 genes that exists in at least two colonies were used for analysis) (Figure 3a).

**Figure 3.**
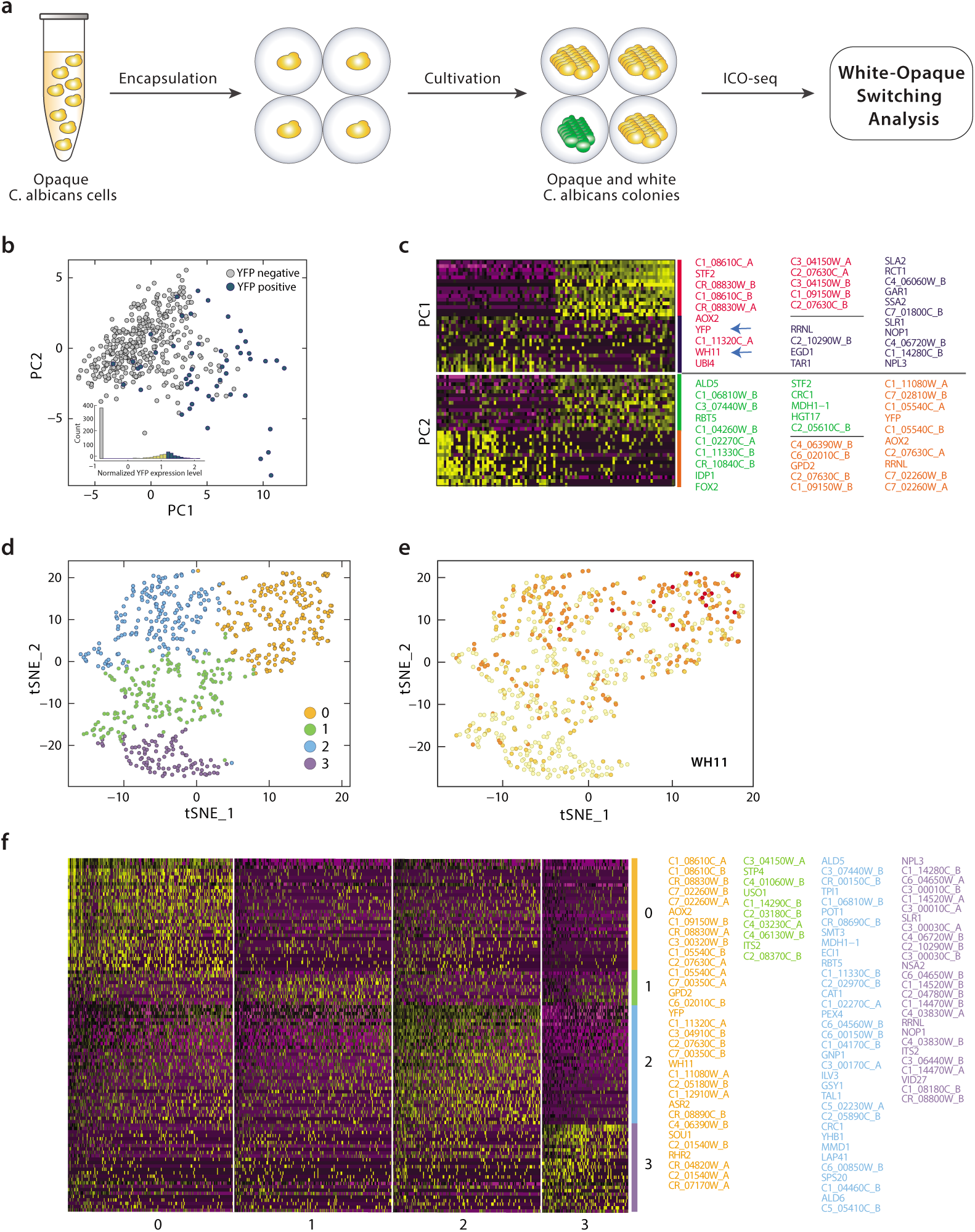
White-opaque switching analysis of *C. albicans* using ICO-seq. (a) Schematic workflow of *C. albicans* opaque-white analysis using ICO-seq. (b) Principal component analysis of gene expression profiles from single *C. albicans* colonies with YFP positive colonies labeled blue and YFP negative colonies labeled grey. (c) Gene expression heatmaps for PC1 and PC2 with both cells and genes are ordered according to their PCA scores (barcodes on x-axis and upregulated gene names on y-axis, yellow indicates upregulation, top 15 genes and 100 barcodes are shown in the heatmap). Genes shown are color coded and listed on the right. (d) t-SNE analysis plot of gene expression profiles from *C. albicans* colonies showing four separate clusters. (e) t-SNE analysis plot with WH11 expression shown in red. (f) Heatmap shows top 35 marker genes (thresh.use set as 0.25) from each cluster. When there were more than 35 genes, only the top 35 were shown. Genes shown are color coded and listed on the right.

Using principle component analysis (PCA) and YFP expression as reference, we saw that colonies with highly positive YFP expression (n=50 with normalized YFP expression > 1.4, which we attribute to white cells), have a distinct gene expression profile from ones with no YFP expression (n=381 with normalized YFP expression < 0, which we attribute to opaque cells) (Figure 3b). Moreover, this PCA analysis revealed that WH11 expression correlates well with YFP expression; in other words, WH11 is also induced in colonies with positive YFP expression (Figure 3c, blue arrow). Finally, PCA analysis revealed that genes implicated in energy metabolism such as C1_08610C, STF2 and AOX2 also correlates with WH11 and YFP. Multiple reports have shown that a majority of the ~1000 gene expression differences (using a two-fold cutoff) between white and opaque cells involve genes implicated in nutrient acquisition and metabolism (31–33). PCA analysis of ICO-Seq data (1) reaffirms that WH11 is a valid reporter for white cells, (2) validates that the major gene expression differences between white and opaque cells involve metabolism and (3) identifies a handful of valuable reference genes for both white and opaque cells.

To gain more insight into the heterogeneity across opaque and white cells with a more unbiased approach, we employed t-SNE (34) using Seurat (35), a clustering algorithm that revealed 4 major clusters (Figure 3d). WH11 expression is mostly contained in colonies within cluster 0 (orange), suggesting cluster 0 corresponds to white colonies (Figure 3e). Besides WH11, genes related to glycerol metabolism and biofilm formation such as GPD2, RHR2 are also upregulated in cluster 0 (Figure 3f), in accordance with previous findings (36,37). In addition, energy metabolism genes (AOX2 and C1_08610C) found by PCA are also over-represented in cluster 0 (Figure 3f). Cluster 2 cells overexpress fermentation genes (ALD5 and 6) (Figure 3f), as is to be expected for opaque cells (38). In addition to these two major populations (which we attribute to white and opaque cells), another cluster (cluster 3, which could either be a separate cell-type or a population of cells that includes both white and opaque cells) is marked by expression of SLR and NPL3 (Figure 3f), members from the SR (serine-arginine) family of RNA-binding proteins that may influence polarized growth (39). This t-SNE analysis uses an orthogonal clustering algorithm to identify additional reference genes that can be used to distinguish between white and opaque cells and further demonstrates the ability of ICO-seq to dissect gene expression differences between cell types.

### Heterogeneity analysis of *S. cerevisiae* ARO4 mutagenesis library

In the previous section, we employed ICO-seq to analyze the differences between two cell-types for which population-level gene expression differences had already been documented. Next, we used ICO-seq to identify gene expression differences across colonies in a fungal system where gene expression profiling has, to our knowledge, not been conducted.

ARO4 is a key regulator for the pathway that synthesizes aromatic amino acids in *S. cerevisiae*, the building blocks of many valuable molecules (40). ARO4 is inhibited by the aromatic amino acid tyrosine, one of several negative feedback loops that constrain flux through the pathway (40). Mutations in the regulatory region (residues 191-263) and elsewhere in the protein have been shown to reduce feedback inhibition by tyrosine and increase metabolic flux through the pathway (6,41). We applied ICO-seq to study the effect that mutations in the regulatory domain of ARO4 have on the transcriptome of *S. cerevisiae* (Figure 4a),

**Figure 4.**
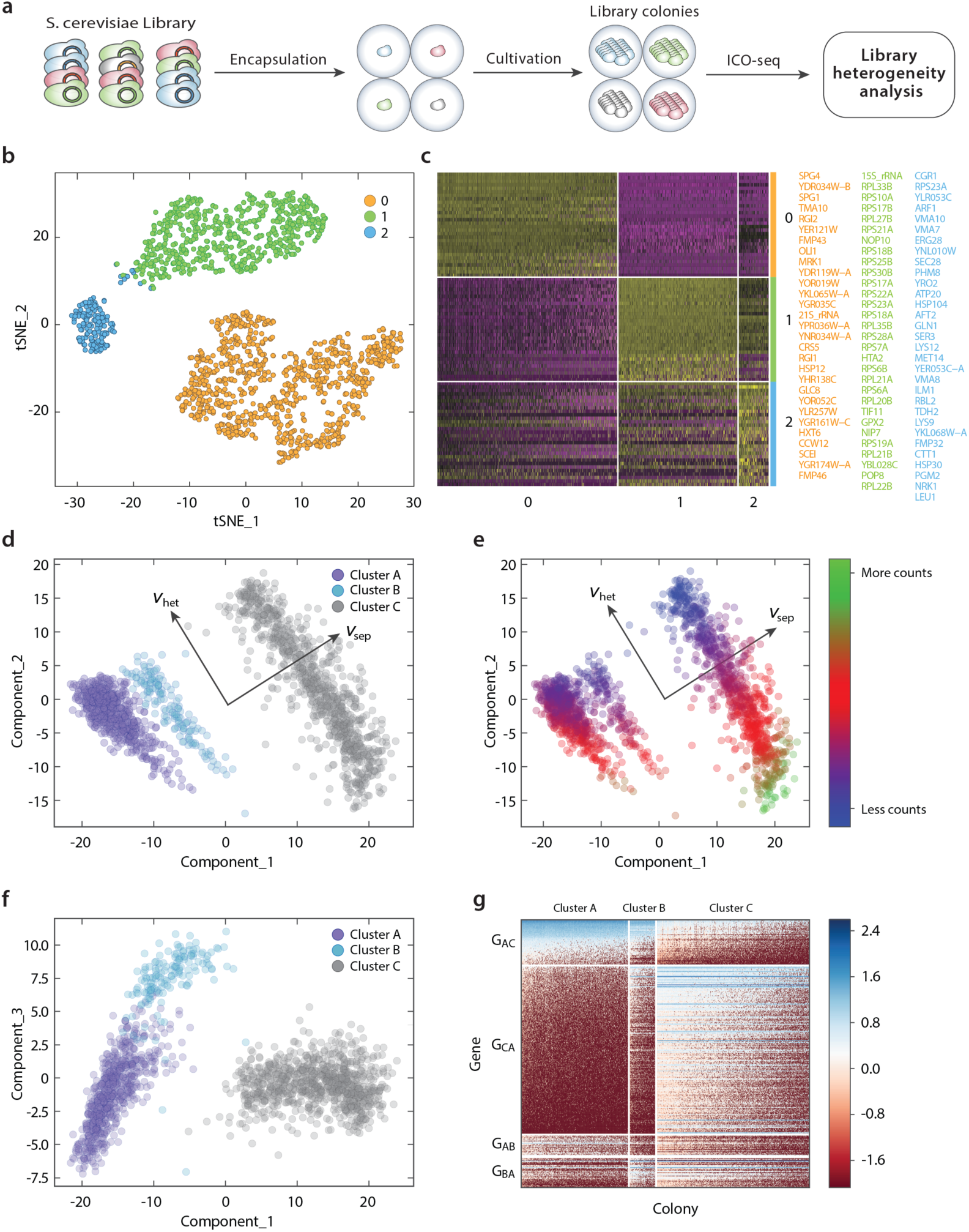
Heterogeneity analysis of S. cerevisiae ARO4 mutagenesis library. (a) Schematic workflow of *S. cerevisiae* library heterogeneity analysis using ICO-seq. (b) t-SNE analysis plot of gene expression profiles from single *S. cerevisiae* colonies showing three clusters. (c) Heatmap shows top 35 marker genes (threshold set as 0.25) from each cluster. When there were more than 35 genes, only the top 35 were shown. Genes shown are color coded and listed on the right. (d) Plot of principal component 1 and 2 values for all colonies analyzed showing the different clusters, as well as the direction that most clearly separates the clusters 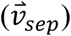 and the direction along which colonies within clusters vary most (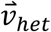). (e) Same as (d), but colonies are colored by the logarithm of the total number of counts. (f) Same as (d) but for principal components 1 and 3. (g) Expression heatmap of genes whose expression separates the clusters most clearly. Groups G_AC_ and G_CA_ include genes that are expressed at higher or lower levels respectively in cluster A than in cluster C. Groups G_AB_ and G_BA_ include genes that are expressed at higher or lower levels respectively in cluster A than in cluster B (subtracting out any genes already included in groups G_AC_ and G_CA_).

We analyzed expression data from 1642 colonies that expressed at least 500 unique genes that exist in at least two colonies. Similar to analysis performed with data from *C. albicans* experiment, we identified three major clusters through tSNE analysis using Seurat (35) (Figure 4b, c). The colonies in cluster 1 express genes related to protein biosynthesis, suggesting these colonies were actively growing when profiled. The colonies in cluster 2 express genes related to amino acid biosynthesis (GLN1, SER3, LYS12, MET14, LYS9, LEU1), suggesting they could correspond to ARO4 mutants that upregulate the amino acid biosynthesis pathway. Meanwhile, colonies in cluster 0 express genes implicated in the stationary phase of growth (SPG4, SPG1) and respiratory metabolism (OLI1, FMP43, RGI2, YDR119W-A).

To provide a more comprehensive analysis, we further limited our analysis to 1064 genes that had at least 3 mRNA counts in at least one of the 300 colonies with the lowest overall expression level. In principal component space we can identify three large clusters that are separated most clearly by a linear combination of principal components 1 and 2 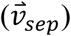 (Figure 4d). Along that axis, clusters A (magenta, N=620) and B (cyan, N=155) express relatively more genes related to cytoplasmic translation (RPL33B, RPS25B, TIF11, RPS18B, etc), and cluster C (grey, N=867) expresses relatively more genes related to growth in stationary phase (SPG1 and SPG4), as well as genes associated with respiratory metabolism (MPC3, OLI1, RGI1, COX26, RGI2) and stress (MRK1, GPX1) (Supplementary Tables 1-3). This suggests that the colonies in clusters A and B were actively growing when profiled. This is further supported by the fact that the correlation between the average expression for clusters A and B to bulk sequencing data from unperturbed cells is higher than the correlation between cluster C and unperturbed bulk sequencing data, while the opposite is true when comparing to bulk sequencing data for cells in which the master regulator PKA is inhibited (Supplementary Figure 1).

There is heterogeneity within the clusters in the direction orthogonal to 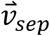(which we denote 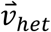). Colonies with lower values on the 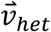axis have higher overall sequencing reads (Figure 4e). These colonies have relatively higher expression of genes associated with the ER-Associated Degradation pathway (UBC7, CUE1) and mitochondrial ribosomal proteins (MRPL38, MRP17, MNP1, MRPS8, MRPL49, Supplementary Table 1). Colonies with lower total transcript counts tend to have higher 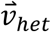values. Those colonies have higher transcript counts of 21s rRNA and RPL24A as well as two paralogous metallothioneins, CUP1-1 and CUP1-2 which mediate resistance to copper and cadmium (Supplementary Table 1). Interestingly, cluster C (which was characterized by higher expression of stress and stationary phase genes) contained the colonies with both the lowest number of transcript counts (which might include colonies that grew very slowly after implanting in a microgel) as well as those with the highest number of transcript counts (which may have slowed growth after reaching saturation).

The two actively growing clusters, A and B, separate most clearly along principal component 3 (Figure 4f, g). Colonies with higher principal component 3 values have relatively more expression of genes associated with the GO term Organophosphate Metabolic Process (YSA1, ATP16, CYC7, PHM8, YNK1, MTD1, PGM2, URA10, NRK1, AFT2) as well as other amino acid metabolism genes (SER3, GLN3, LYS12) and relatively lower expression of genes associated with the GO term rRNA processing (POP8, NHP2, LOC1, CGR1, LRP1, EBP2, CBF5, RPL37A, NOP56, NOP15, NIP7, Supplementary Tables 1-3). This suggests that the colonies in cluster B could be actively growing mutants from the ARO4 library that have adjusted their metabolism to adapt to changes in metabolic flux resulting from the mutations.

From this analysis, we can group colonies from a yeast mutagenesis library into three major bins and use gene expression profiling to select those that appear to be conducting efficient protein biosynthesis. Confirming that the observed heterogeneity is a direct result from the mutations of ARO4 will require further development of ICO-seq to enable the direct linking of genotype (mutations) with phenotype (transcriptome). We expect that using high-throughput RNA sequencing to screen for mutants with desired gene expression profiles will be valuable for protein engineering (42), deep mutational scanning (43), and strain engineering (44); Current methods are costly, limited in throughput, or restricted to optical assays (6,8). ICO-seq may provide a versatile, high-throughput and scalable alternative for such analyses going forward.

## Discussion

ICO-seq enables high-throughput RNA-seq of isogenic cell colonies, which can be used to characterize cellular phenotype for high throughput screening applications. We demonstrated the proof-of-concept usage of ICO-seq by expression profiling hundreds to thousands of colonies of both *C. albicans* and *S. cerevisiae*. By scaling up our protocol, we would be able to collect data from hundreds of thousands of individual yeast colonies. Key to the method is the capture, growth, and sequencing of single cells in agarose hydrogel spheres, which affords a facile route towards generating and analyzing large numbers of distinct colonies. Because gene expression is a universal readout, ICO-seq can be applied to a broad range of phenotypes such as metabolic flux prediction (45–47). It can also be applied to dissect a heterogeneous response from single colonies when they are perturbed (e.g. addition of toxic compound, elevated temperature and when co-cultured with other species). Moreover, microgels are compatible with sorting via flow cytometry prior to ICO-seq, adding more flexibility of the workflow. Additionally, collective analysis of gene expression across large numbers of genetically distinct variants can be applied to characterize sequence-phenotype relationships, and aid in identifying optimal genetic engineering to obtain a desired phenotype. To do this, it is essential to link gene expression profiles to perturbation genotypes. This can be done by incorporating barcodes (expressed as mRNA) for each perturbation as is done in Perturb-seq applications (48,49). In this method a library of single-guide RNA (sgRNA) that cause CAS9 based genetic perturbations are introduced into single cells and transcriptome data is collected using droplet-based sequencing. The identity of the sgRNA for each perturbation is encoded in a sequence that is highly expressed in each cell and is sequenced with the rest of the transcriptome. ICO-seq will enable Pertub-seq to be used with yeast and potentially other microbes.ICO-seq can also be applied to genetic circuits, allowing characterization of genetic logic gates and associated promoters, insulators, and terminators, for thousands of circuit variants per experiment (50).

While we have focused on yeast, ICO-seq is applicable to any culturable cell type, including mammalian, bacterial, plant and other fungal cells. Profiling soil microbes may yield novel enzymes and pathways (51), and analyzing the gene expression of gut microbes when cultured in the gut may yield insights useful to microbiome-based therapy (52). By combining ICO-seq with modern *in situ* cultivation approaches (53) that also rely on hydrogel colony culture, it should be possible to obtain accurate and comprehensive gene expression profiles for currently uncultivable organisms. When applied to mammalian cells, ICO-seq should be valuable for methods that currently rely on single cell RNA-seq, such as Perturb-seq (48,49), with the significant benefit of reduced expression noise. ICO-seq also enables new possibilities for studying small colonies of genetically or phenotypically distinct cells, allowing accurate gene expression profiling of microbial consortia, cancer spheroids (54) or organoids (55,56). While we have applied the approach using a custom microfluidics and Drop-seq strategy, new microgel workflows announced for commercial platforms (10X Genomics) should allow our approach to be applied in a more accessible format using commercially available hardware.

## Methods

### Microfabrication of devices

Photoresist masters are created by spinning on a layer of photoresist SU-8 (Microchem) onto a 3 inch silicon wafer (University Wafer), then baking at 95 °C for 20 min. Then, the photoresist is subjected to 3 min ultravoilet exposure over photolithography masks (CAD/Art Services) printed at 12,000 DPI. After ultravoilet exposure, the wafers are baked at 95 °C for 10 min then developed in fresh propylene glycol monomethyl ether acetate (Sigma Aldrich) then rinsed with fresh propylene glycol monomethyl ether acetate and baked at 95 °C for 5 min to remove solvent. The microfluidic devices are fabricated by curing poly(dimethylsiloxane) (10:1 polymer-to-crosslinker ratio) over the photoresist master (57). The devices are cured in an 80 °C oven for 1 h, extracted with a scalpel, and inlet ports added using a 0.75 mm biopsy core (World Precision Instruments). The device is bonded to a glass slide using O_2_ plasma treatment and channels are treated with Aquapel (PPG Industries) to render them hydrophobic. Finally, the devices are baked at 65 °C for 20 min to dry the Aquapel before they are ready for use.

### Yeast strains and culture conditions

*C. albicans* experiments were carried out in strain RZY122. RZY122 is modified from strain SN87, a SC5314 derivative (58), with YFP-HIS1 replaces one copy of the WH11 ORF (a/a leu2Δ/leu2Δ his1Δ/his1Δ URA3/ura3Δ::imm IRO1/iro1Δ::imm WH11::YFP-HIS1/WH11). Cultures of opaque cells were inoculated from a single colony from a plate into liquid media at 25 °C for ~20 hours before encapsulation. Plates are standard synthetic complete yeast media with Uridine. Media is same as the plates but without agarose.

*S. cerevisiae* experiments were carried out in strain BY4741 (MATa; his3Δ1; leu2Δ0; met15Δ0; ura3Δ0). The ARO4 regulatory domain mutagenesis library used in this study has been described in Abatemarco et al. (6). The library was cultivated with constant agitation in YSC-HIS at 30 °C. YSC-HIS media consisted of 20 g/L glucose (Thermo Fisher Scientific), 0.77 g/L CSM-His supplement (MP Biomedicals), and 0.67 g/L Yeast Nitrogen Base (Becton, Dickinson, and Company).

### Agarose microgel generation

The yeast culture was washed with PBS 2 times before being resuspended in PBS at an appropriate concentration based on hemocytometer counting. The low melting agarose (Sigma) solution was made with heating 2% ultra low melting agarose in PBS at 90 °C until completely melted. The melted agarose was quickly loaded in a syringe and installed to the pump. A tabletop space heater set to 80 °C was positioned to keep the agarose syringe warm during droplet generation. HFE7500 oil with 2% ionic krytox as surfactant was used for the oil phase. The agarose droplets were collected into a 50 ml falcon tube placed on ice for the formation of agarose microgel. The agarose microgel were released from the droplets by adding 20% PFO (1H,1H,2H,2H-Perfluoro-1-octanol, sigma) in HFE7500 into the emulsion followed by washing twice with TETW solution (10mM Tris pH 8.0, 1mM EDTA, 0.01% Tween-20). The agarose microgels were then resuspended in appropriate media for overnight culturing to form single colony containing microgels.

### Dropseq workflow with agarose microgels

Barcoded Drop-seq beads were purchased from ChemGenes corporation (catalog number MACOSKO-2011-10) at 10um synthesis scale. Cleanup and storage of the beads was performed as in (17).

For the *C. albicans* experiment, the barcoded Drop-seq beads were resuspended in 0.9X YR lysis buffer (Zymo Research) with additional 500 mM NaCl. The yeast colonies in agarose microgels were first washed with PBS buffer twice and YR Digestion buffer (Zymo Research) once then treated with Zymolyase in YR Digestion buffer to digest the yeast cell walls for 1 hour at 37°C. The microgels were then close-packed in a 1ml syringe by centrifugation. The dropseq beads, close-packed agarose microgels, and HFE 7500 with 2% w/v EA surfactant (RAN Biotechnologies) were then loaded into the yeast agarose Drop-seq device for droplet generation controlled via syringe pumps.

For the *S. cerevisiae* experiment, a LiOAc-SDS based lysis protocol was used to simplify the workflow (59). The barcoded Drop-seq beads were resuspended in Drop-seq lysis buffer with additional 400 mM LiOAc, 2% SDS solution and 500mM NaCl. The microgels were then close-packed in a 1ml syringe by centrifugation. The dropseq beads, close-packed microgels, and HFE 7500 with 2% w/v ionic krytox surfactant were then connected to the yeast agarose Drop-seq device for droplet generation. The collected emulsions were heated at 70 °C for 15min and then kept on ice to facilitate cell lysis and mRNA capture.

The collected emulsions were then processed following the Drop-seq protocol (17). In short, collected emulsions were broken by addition of PFO. mRNA transcripts bound on Drop-seq beads were then reverse transcribed using reverse transcriptase (Maxima RT, Thermo Fisher). Unused primers were degraded by addition of Exonuclease I (New England Biolabs). Washed beads were counted and aliquoted for PCR amplification. 600 pg of the amplified cDNA was then used to construct a sequencing library using the Nextera XT kit.

### Next generation sequencing and data analysis

Nextera XT generated sequencing libraries were sequenced with Illumina MiSeq system using custom read 1 primer. Read1 length is 25 bp and Read2 length is 75 bp. *C. albicans* library was sequenced with 24 million paired-end reads and *S. cerevisiae* library was sequenced with 23.6 million paired-end reads.

The sequencing reads were fed into the Drop-seq bioinformatics analysis pipeline following Drop-seq protocol (17) except the alignment were done with Bowtie (60). *C. albicans* (61) or *S. cereivisae* (62) reference genomes were used for alignment. YFP gene sequence was added to the *C. albicans* reference genome to facilitate the analysis. The resulted digital gene expression tables were then used as input for Seurat V1.3 for analysis and visualization (35). For *C. albicans* PCA plot, we first used scale.data from Seurat object to analyze the expression distribution of YFP gene followed by extracting cell barcodes with high YFP expression (>1.4) and low YFP expression (<0) to use as white cells and opaque cells respectively. We scanned different cut-offs and parameters to balance data quality and cell numbers.

For the Principal Component Analysis for the ARO4 library, we first filtered out genes that did not have at least 3 counts in at least one of the 300 colonies with the lowest overall expression level. We then normalized counts for each colony, *k* by the total number of read counts for that colony in this reduced set of genes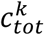 and multiplied by the median total read number for all colonie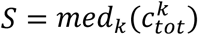. We then added a pseudocount of 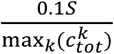 so that the pseudocount would be 10% of the value of the smallest possible value that a real count could take on. We then took the base 10 logarithm prior to principal component analysis (Figure 4c-f). The vector 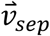 is defined as the direction connecting the points(*x*_0_, *y*_0_)= (3.1, −1.6) and(*x*_1_, *y*_1_)= (20.9, 10.5) in PC1 PC2 space. The vector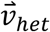 is the vector perpendicular to 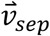 that passes through *x*_0_, *y*_0_ and *x*_2_, *y*_2_ = 3(−6.9, *y*_2_). For each colony we can find coordinates in PC1/PC2 space, and project those coordinates onto 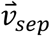 to obtain *u*_*sep*_Clusters A, B, and C are defined as all colonies for which *u*_*sep*_< −9.83, −9.83 < *u*_*sep*_< 0, or *u*_*sep*_> 0 respectively.

## Data availability

The data that support the findings of this study are available from the corresponding author upon reasonable request.

## Acknowledgements

We thank Dr. Alexander D. Johnson from UCSF for gifting *C. albicans* strain RZY122. We also thank Dr. Alexander Johnson, Dr. John Haliburton, and Dr. Cyrus Modavi for helpful discussion. This work was supported by the Chan Zuckerberg Biohub, the National Science Foundation CAREER Award (Grant Number DBI-1253293); the National Institutes of Health (NIH) (Grant Numbers R01-EB019453-01, R01-HG008978 and DP2-AR068129-01.) Chiraj Dalal was supported by NIH grant R01 AI049187 to AJ.

### Competing Interests

The authors declare that they have no competing financial interests.

### Correspondence

Correspondence and requests for material should be addressed to A.R.A.

### Contribution

A.R.A and L.L. conceived the project. L.L. C.D. and B.H. designed and carried out the experiments. L.L., C.D. and B.H. and A.R.A. evaluated the data. L.L. and A.R.A. wrote the manuscript with input from C.D. and B.H..

